# Rigidity of silicone substrates controls cell spreading and stem cell differentiation

**DOI:** 10.1101/040048

**Authors:** Grigory Vertelov, Edgar Gutierrez, Sin-Ae Lee, Edward Ronan, Alex Groisman, Eugene Tkachenko

## Abstract

Multiple functions of cells cultured on flat substrates have been shown to depend on the elastic modulus of the substrate, E, with the dependence being strongest in a physiological range of soft tissues, corresponding to E from 0.1 to 100 kPa. Among those functions are stem cell differentiation, cell spreading, and cell signaling [1]. In the context of differentiation of mesenchymal stem cells (MSCs), substrates with E in the ranges of <4 kPa, 8-17 kPa, and >25 kPa, have been classified as soft (adipogenic) [2,3], medium rigidity (myogenic)1, and hard (osteogenic) [1], respectively. In most studies, the soft substrates are hydrogels, and variations in their elastic moduli are usually accompanied by variations in the dry mass and porosity. The paradigm of the effect of substrate rigidity on the cellular functions has been challenged by Trappmann et al. [4], who claimed that cell spreading and differentiation on hydrogel substrates depend not on the elastic moduli of the substrates, but rather on their porosity, which affects the density of adhesion points between the substrate surface and the extracellular matrix (ECM) coating on it. This claim has been rebutted by Wen at al. [3], who have used hydrogel substrates with different porosities but identical elastic moduli to show that it is the elastic modulus rather than the porosity that is key to the effect of the substrate on cell spreading and differentiation. Both publications agree, however, that there is no appreciable effect of the substrate rigidity on either cell spreading or differentiation, if the substrate is made of a silicone gel rather than a hydrogel. This conclusion appears to contradict the findings of several other groups, who reported that when cells are plated on an array of flexible silicone microposts, their spreading and differentiation depend on the rigidity of the substrate [5], and that when cell are plated on silicone gels, their differentiation depends on the gel rigidity [6]. To resolve this contradiction, we used soft, medium, and hard silicone gel substrates with elastic moduli of 0.5, 16, and 64 kPa, respectively, (Fig.1) to perform experiments similar to those reported in Refs.4 and 3, testing the dependence of differentiation and spreading of MSCs and of spreading of fibroblasts and keratinocytes on the substrate rigidity.

---

Multiple functions of cells cultured on flat substrates have been shown to depend on the elastic modulus of the substrate, *E*, with the dependence being strongest in a physiological range of soft tissues, corresponding to *E* from 0.1 to 100 kPa. Among those functions are stem cell differentiation, cell spreading, and cell signaling^1^. In the context of differentiation of mesenchymal stem cells (MSCs), substrates with *E* in the ranges of <4 kPa, 8-17 kPa, and >25 kPa, have been classified as soft (adipogenic)^2,3^, medium rigidity (myogenic)^1^, and hard (osteogenic)^1^, respectively. In most studies, the soft substrates are hydrogels, and variations in their elastic moduli are usually accompanied by variations in the dry mass and porosity. The paradigm of the effect of substrate rigidity on the cellular functions has been challenged by *Trappmann et al.*^4^, who claimed that cell spreading and differentiation on hydrogel substrates depend not on the elastic moduli of the substrates, but rather on their porosity, which affects the density of adhesion points between the substrate surface and the extracellular matrix (ECM) coating on it. This claim has been rebutted by *Wen at al.*^3^, who have used hydrogel substrates with different porosities but identical elastic moduli to show that it is the elastic modulus rather than the porosity that is key to the effect of the substrate on cell spreading and differentiation. Both publications agree, however, that there is no appreciable effect of the substrate rigidity on either cell spreading or differentiation, if the substrate is made of a silicone gel rather than a hydrogel. This conclusion appears to contradict the findings of several other groups, who reported that when cells are plated on an array of flexible silicone microposts, their spreading and differentiation depend on the rigidity of the substrate^5^, and that when cell are plated on silicone gels, their differentiation depends on the gel rigidity^6^. To resolve this contradiction, we used soft, medium, and hard silicone gel substrates with elastic moduli of 0.5, 16, and 64 kPa, respectively, (Fig.1) to perform experiments similar to those reported in Refs.^4^ and ^3^, testing the dependence of differentiation and spreading of MSCs and of spreading of fibroblasts and keratinocytes on the substrate rigidity.

**Figure 1.**
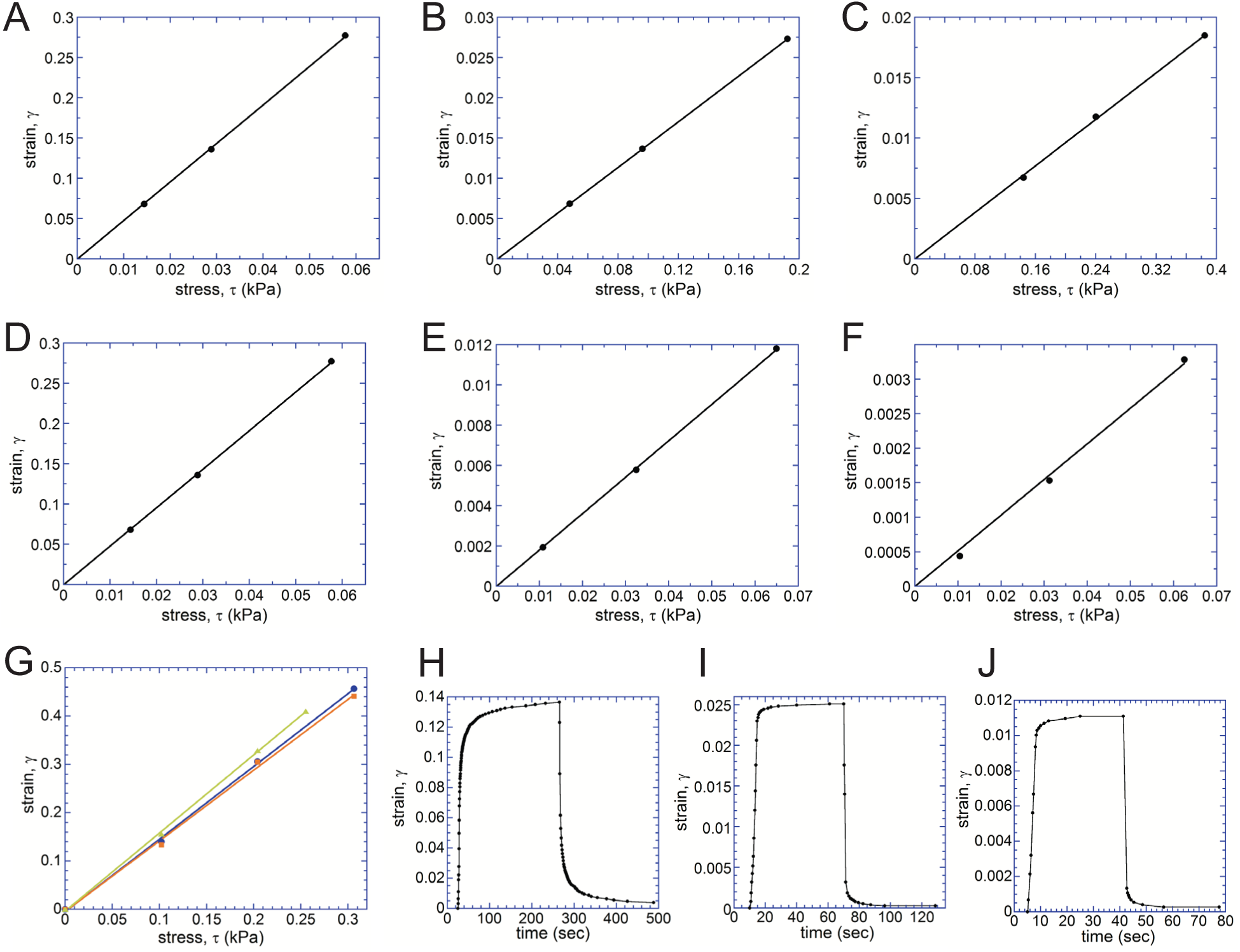
Measurements of the elastic moduli of the silicone gels. Elastic moduli of the silicone gels used in the experiments on cells were tested by measuring the deformation of thin uniform layers of gels and calculating their shear strain, γ, under known shear stresses, τ, using a previously reported microfluidic technique^1^ (A) - (C) and a specially built gel rheometer^2^ (D) - (F). (A), (B), and (C) Dependence of γ on τ for gels with nominal elastic moduli of 0.5, 16, and 64 kPa, respectively. The thicknesses of the gel layers are 37, 31, and 63 μm, respectively (special order from MuWells). All three dependencies are fitted by straight lines passing through the origin, and the elastic moduli of the gels are calculated from the fits as *E* = (3*γ* / *τ*)^−1^ at 0.61, 20, and 62 kPa for the 0.5, 16, and 64 kPa gels respectively. (D), (E), and (F) Dependence of γ on τ for gels with nominal elastic moduli of 0.5, 16, and 64 kPa, respectively. The thicknesses of the gel layers are 1.40, 1.35, and 1.54 mm, respectively (special order from MuWells). All three dependencies are fitted by straight lines passing through the origin, and the elastic moduli of the gels are calculated from the fits as *E* = (3*γ* /*τ*)^−1^ at 0.40, 17, and 65 kPa for the 0.5, 16, and 64 kPa gels, respectively. (G) Dependence of γ on τ for a silicone gel with a nominal elastic modulus of 2 kPa was measured for gel layers (all special orders from MuWells) with thicknesses of 18.2 μm (*blue circles*), 6.1 μm (*orange squares*), and 2.40 μm (*green triangles*) using a modified microfluidic technique with Helium as the working fluid. All three dependencies are fitted by straight lines passing through the origin, and the elastic moduli of the gel layers are calculated from the fits as *E* = (3*γ* / *τ*)^−1^ at 1.70, 1.78, and 1.65 kPa for the 18.2, 6.1, and 2.40 μm thick gel layers, respectively. A measurement with a shear rheometer on a 1.05 mm thick gel layer resulted in an elastic modulus of 1.90 kPa. (H) - (J) Incremental shear strains, γ, of the 0.5, 16, and 64 kPa SoftSubstrates gels as functions of time under step-wise changes of shear stress, τ. The measurements were performed with the shear rheometer on 0.5, 16, and 64 kPa gels gel layers with respective thicknesses of 0.43, 2.55, and 2.52 mm (all special orders from MuWells; the actual values of *E* were measured at 0.44, 18, and 65 kPa, respectively). For the 0.5 kPa gel (H), the value of τ was increased from 0.42 to 20.6 Pa at 25 sec and reduced back to 0.42 Pa at 265 sec; for the 16 kPa gel (I), the value of τ was increased from 2.5 to 160 Pa at 10 sec and reduced back to 2.5 kPa at 70 sec; for the 64 kPa gel (J), the value of τ was increased from 201 to 487 Pa at 10 sec and reduced back to 201 Pa at 70 sec. The transition curves for γ after changes in τ were not well-fitted by single exponentials, which was not surprising, because a gel usually has a broad spectrum of relaxation times. Therefore, as representative relaxation times, we took times it took the transition to be 63% (1-e^−1^) complete. For the reductions in τ, the relaxation times were 3 sec for the 0.5 kPa gel and <1 sec for both 16 and 64 kPa gels. The temporal resolution of the measurement technique is on the order of 1 sec. Therefore, the relaxation times measured for the 16 and 64 kPa gels were experimentally indistinguishable from zero. For the increases in τ, the relaxation times were 5.5 sec for 0.5 kPa, 2.5 sec for 16 kPa, and 1.5 sec for 64 kPa. The rheometer operates in such a way, however, that it takes ~2 sec for τ to increase (a reduction in τ occurs faster). Therefore, the gel relaxation times obtained from the step-wise increases and decreases of τ are consistent, and when corrected for the instrumental delays and time resolution, the relaxation times of the 0.5, 16, and 64 kPa gels can be estimated at <4, <1, and <1 sec, respectively.

In an adipogenic medium (Fig.2,3), when MSCs were plated on the 64 kPa substrate, their differentiation to adipocytes somewhat increased as compared to a plastic substrate control, and when the MSCs were plated on the 16 kPa and 0.5 kPa substrates, their differentiation to adipocytes increased >3-fold. In an osteogenic medium (Fig.2,3), the differentiation of MSCs to osteoblasts was reduced to ~80% on the 64 kPa substrates as compared with a plastic control and was further reduced to ~36% on the 16 kPa substrates and to ~27% on the 0.5 kPa substrates, with the differences between the three substrates and the control being all significant. The average spreading areas of MSCs were significantly smaller on the 0.5 kPa silicone gel than on the 16 and 64 kPa gels (Fig.4). The average areas of both keratinocytes and fibroblasts cultured on the silicone gel substrates monotonically increased with the substrate elastic moduli, with differences in the cell areas between the three substrate rigidities being all significant for both cell types (Fig.5). In agreement with the previous report^7^, we found the phosphorylation level of FAK to monotonically increase with the substrate rigidity for both keratinocytes and fibroblasts (Fig.5). Finally, deformations of the silicone gel substrates by traction forces of adherent fibroblasts were inverse functions of the substrate rigidity and had magnitudes comparable to those reported on hydrogels of similar elastic moduli^3,8^ (Fig.6). Therefore, in all four types of assays, the dependence of the cellular functions on the substrate rigidity was qualitatively the same as for cells cultured on hydrogels and micropost arrays, suggesting that the effects of substrate rigidity on functions of plated cells are similar for all types of deformable substrates.

**Figure 2.**
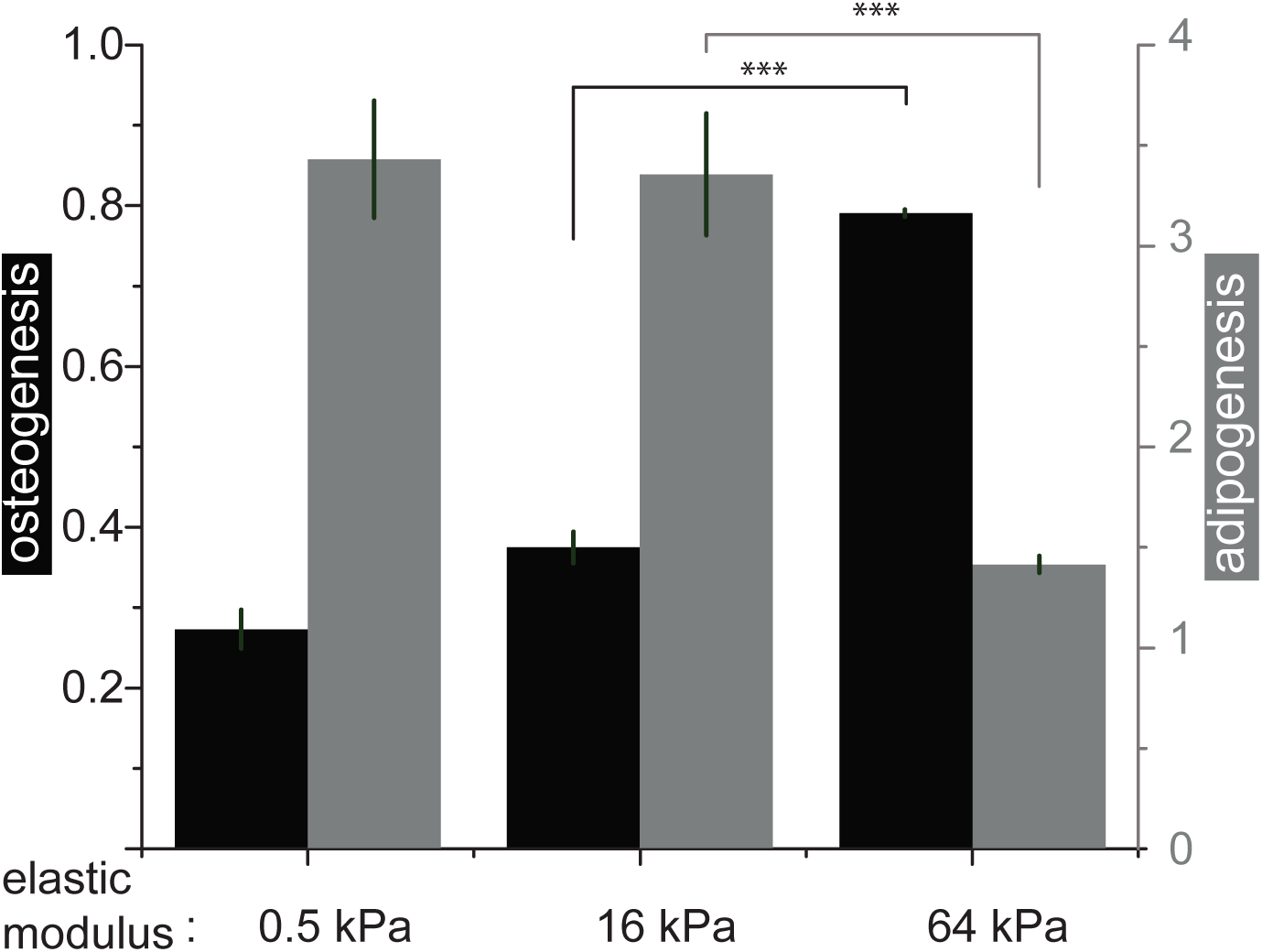
Differentiation of stem cells on silicone substrates with different rigidities. Chemically induced osteogenesis (*black*) and adipogenesis (*grey*) of human mesenchymal stem cells (hMSCs). *Left* and *right ordinates* indicate the levels of differentiation to osteoblasts and adipocytes, respectively, (n=3 wells; representative results from 3 independent experiments) normalized to the levels of differentiation of hMSCs plated on plastic surfaces. *** - p<0.01.

**Figure 3.**
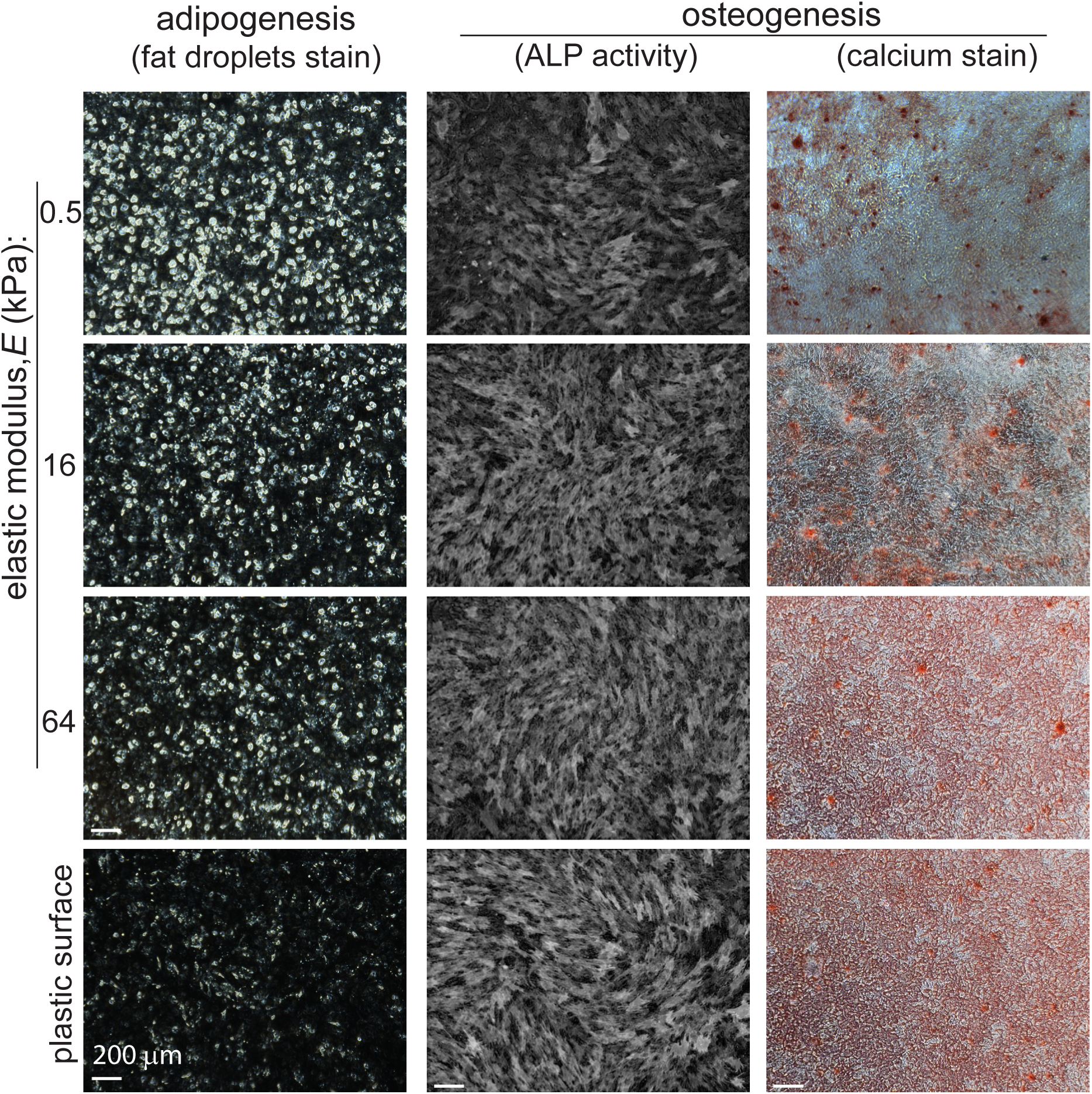
Differentiation of stem cells on substrates of different rigidities. Chemically induced adipogenesis (*left column*) and osteogenesis (*two right columns*) of hMSCs cultured on silicone gel substrates with different elastic moduli. The *brightness* in the *left column* shows fluorescent staining of adipocytes after 14 days of differentiation. The *brightness* in the *middle column* illustrates alkaline phosphatase activity staining after 7 days of differentiation. *Red color* in the *right column* corresponds to calcium staining after 14 days of differentiation.

**Figure 4.**
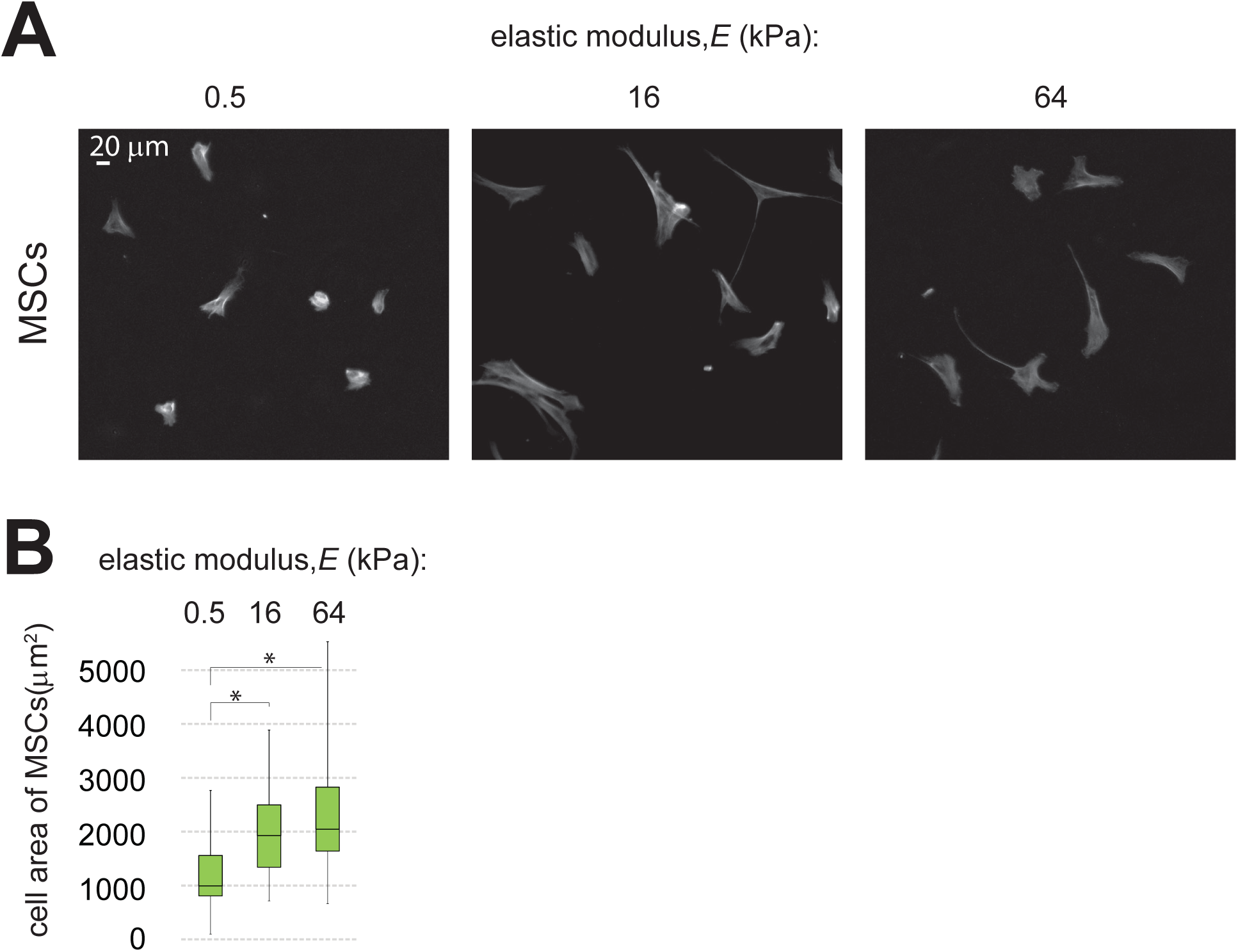
Spreading of stem cells on substrates of different rigidities. (A) Representative fluorescence images of hMSCs on silicone substrates with elastic moduli of 0.5 16, and 64 kPa. The substrates were coated with collagen I and cells were stained with phal-loidin to fluorescently label F-actin. (B) Spreading areas of hMSCs on silicone substrates with different elastic moduli obtained from the analysis of the fluorescence images. Box corresponds to interquartile range of cell spreading areas; black line indicates median value; whiskers show minimal and maximal values. N=40 cells for each type of substrates. *- statistical significance with p<0.01.

**Figure 5.**
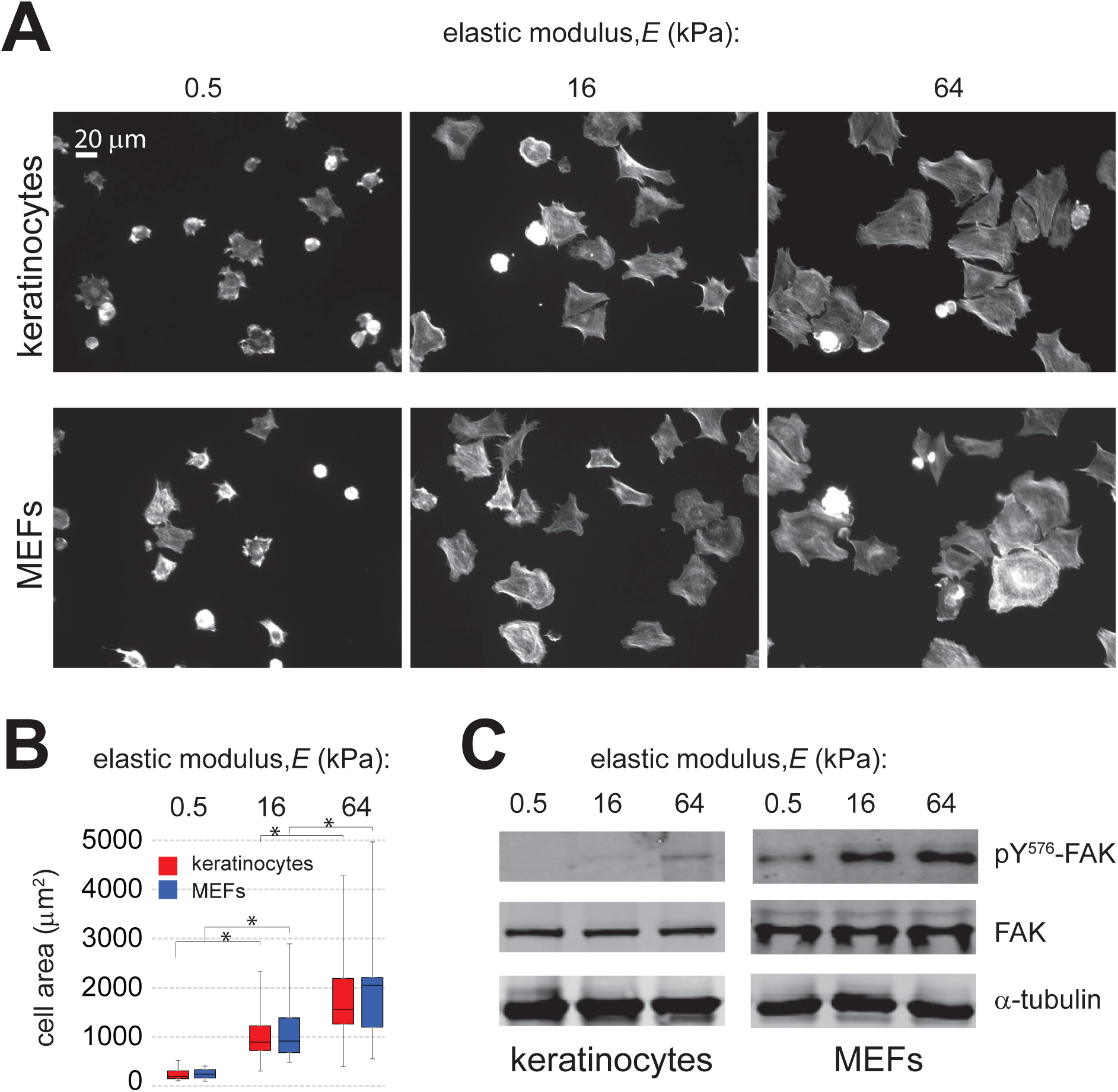
Spreading and signaling of cells on substrates of different rigidities. (A) Representative fluorescence images of keratinocytes and fibroblasts on silicone substrates with elastic moduli of 0.5 16, and 64 kPa. The substrates were coated with fibronectin and cells were stained with phalloidin to fluorescently label F-actin. (B) Spreading areas of keratinocytes (*red*) and fibroblasts (*blue*) on silicone substrates with different elastic moduli obtained from the analysis of the fluorescence images. Box corresponds to interquartile range of cell spreading areas; black line indicates median value; whiskers show minimal and maximal values. N=75 cells for each cell type on substrates of each elastic modulus. *- statistical significance with p<0.01. (C) The levels of phosphorylation of FAK at Tyr-576 for keratinocytes and fibroblasts plated on substrates of different elastic moduli (*upper panels*). *Middle panels* show the total amount of FAK, and *low panels* indicate the amount of α-tubulin, a gel loading control.

**Figure 6.**
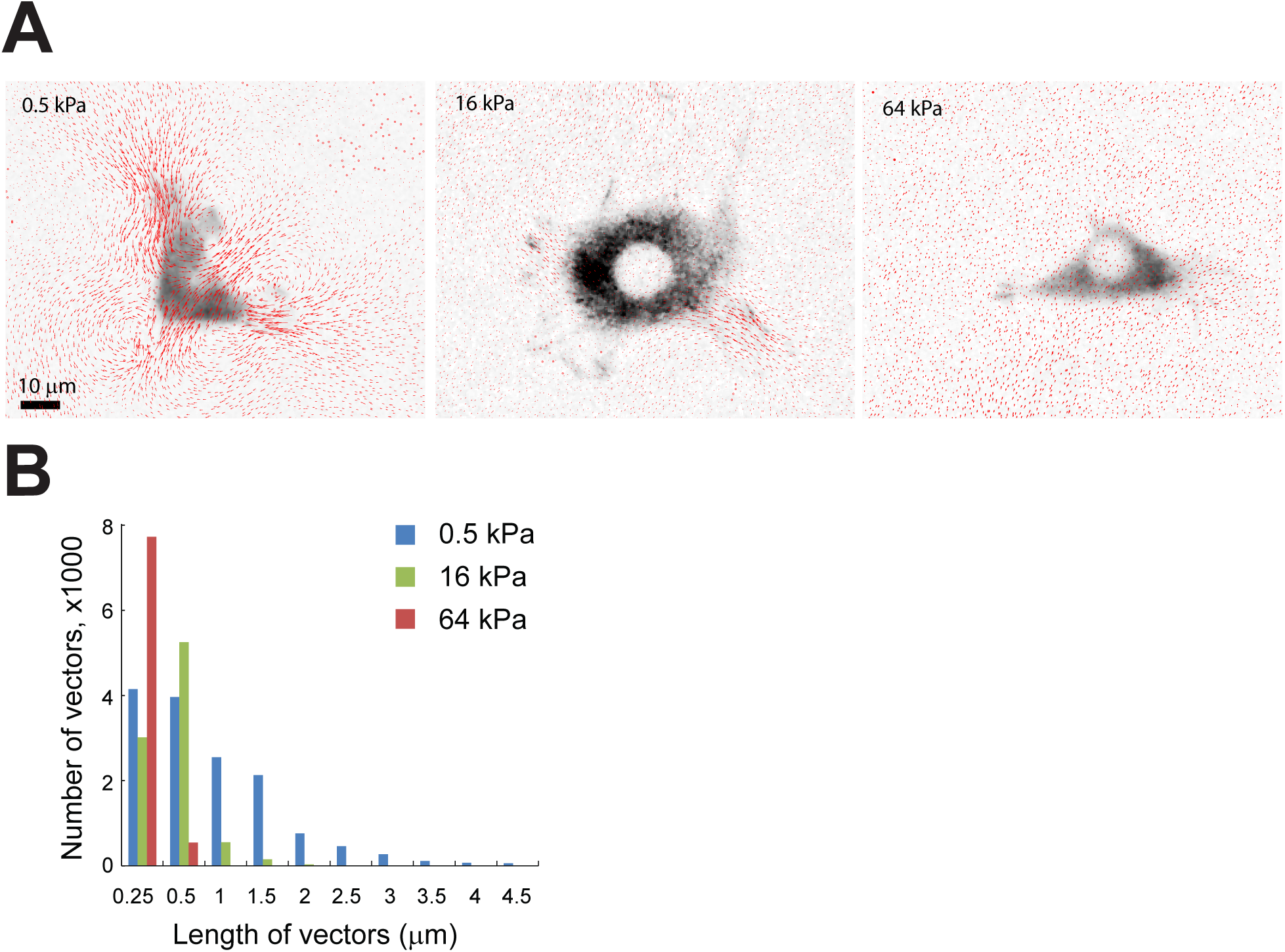
Cell-induced deformations of substrates of different rigidities. (A) Inverted greyscale fluorescence images of fibroblasts plated on substrates with *E* = 0.5, 16, and 64 kPa superimposed with vector maps of the surface displacement (*red arrows*). Fibro-blasts are expressing paxillin-mCherry. Vector maps are obtained by tracking the displacements of 40nm red fluorescent beads attached to the substrate surface.^3,9^ (B) Histograms of the lengths of surface displacement vectors for substrates of different rigidities.

To explain the discrepancies between our findings and the conclusions of Refs.^4^ and ^3^ we note that, whereas we plated cells on substrates from all three ranges of rigidity, none of the silicone gel substrates used in Refs.^4^ or ^3^ was clearly shown to be either soft or of medium rigidity (Fig.7). For hard gel substrates (*E* ≥ 40 kPa), one can expect the dependence of cellular functions on the substrate elastic modulus to be weak. In addition, the surfaces of silicone gel substrates used in our study have amino-reactive groups (Fig.8), providing covalent binding of ECM proteins similar to the binding of ECM to the surfaces of hydrogels in Refs.^4^ and ^3^. It is not completely clear, whether the ECM binding to the silicone gel surfaces used in Refs.^4^ and ^3^ was covalent or passive, and as argued in both papers, cellular responses to the substrate rigidity are expected to depend on the details of binding of ECM to the substrate.

**Figure 7.**
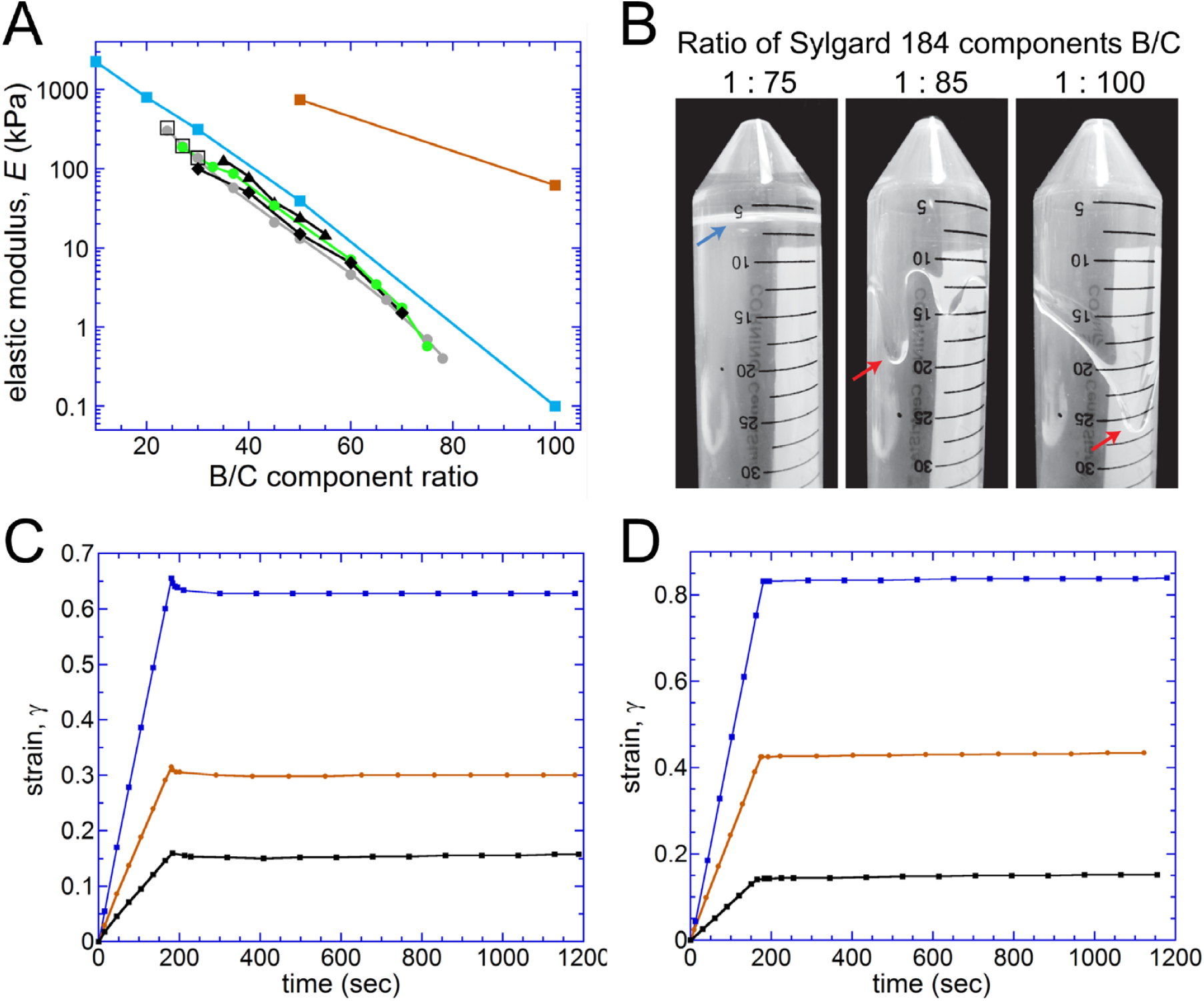
Mechanical properties of materials obtained from mixing different proportions of the base (B) and cross-linker (C) components of Sylgard 184 elastomer. (A) Elastic moduli, *E*, of different mixtures of the B and C components of Sylgard 184 after curing, as measured by several groups using different techniques. *Black triangles* (and *black line*): measurements of axial stress of cylindrical samples as a function of the applied axial extension, from Cesa *et al.*^7^ *Black diamonds:* measurements of axial stress of cylindrical samples as a function of the applied axial extension, from Schellenberg *et al.*^12^. *Open squares*: measurements of extension as a function of longitudinal stress applied to a slab, from Gutierrez *et al.*^1^. *Grey circles* (and *grey line*): measurements of shear strain of ~ 70 μm layer of gel as a function of hydrodynamic shear stress in a microfluidic setup, from Gutierrez *et al.*^1^ *Green circles* (and connecting *green line*): measurements of shear strain as a function of shear stress with a specially built rheometer ^2^. We performed the measurements over the course of two years on samples prepared from different batches of Sylgard 184. The samples were 1-2 mm thick and had lateral dimensions of 20x30 mm. *Blue squares* (and connecting *blue line*): measurements using indentation of gel layers, from Trappmann *et al.*^5^. *Red squares* (and connecting *red line*): values of *E* estimated based on the data from Fig. 4d in Wen *et al.*^6^ by comparing the apparent stiffness of Sylgard silicone samples with that of polyacrylamide (PAA) gels of known elastic moduli (1 and 30 kPa). Wen *et al.*^6^ quantified the stiffness as a substrate spring constant by measuring the resistance of the samples to the retraction of a probe of an atomic force microscope (AFM). For the two PAA gels at low speeds of the probe retraction, the ratio between the apparent substrate spring constant (in pN/mm) and *E* (in kPa) averaged at ~6.25. This average ratio was used to convert the apparent spring constants measured at low probe retraction speeds for silicone samples with B:C = 50:1 and 100:1, ~120 and ~10 pN/mm, respectively, into elastic moduli, *E* = 750 and 63 kPa, respectively. (B) The base and curing agent components of silicone elastomer Sylgard 184 (Dow Corning) were mixed at ratios of B:C = 75:1, 85:1, and 100:1, in conical 50 ml tubes using an overhead stirrer. The silicone pre-polymer was spun down by centrifugation, and the test tubes were baked overnight in a forced convection oven at 80 °C. The test tubes were removed from the oven, allowed to cool to room temperature, turned upside down for 20 min and then photographed. *Blue arrow* points to the flat surface of solidified silicone gel, which was formed from the 75:1 mixture. *Red arrows* point to the surfaces of the silicone in the test tubes with the 85:1 and 100:1 mixtures. Uneven contours of the silicone surfaces in the last two test tubes show that both materials flow under gravity and are thus fluids rather than solids. We have tested different batches of Sylgard 184 elastomer multiple times, mixing its components at different proportions, baked the mixtures at temperatures up to 100 °C for multiple days, but have never been able to make mixtures with B:C ≥ 80:1 produce solid materials. C) Mechanical tests of the material obtained by mixing the components of Sylgard 184 at B:C = 85:1 and curing the mixture overnight in a forced convection oven at 80 °C. Using the shear rheo-emeter, at time t = 0 a 184 μm thick layer of the material was subjected to a certain shear stress, at a time ~180 sec the shear stress was dropped to zero, and the shear strain of the layer was measured as a function of time. The values of the shear stress applied between 0 and 180 sec were 4.68 Pa (black dots), 9.16 Pa (orange dots), and 18.7 Pa (blue dots). The segments with the constant non-zero shear stress are well fitted by straight lines with slopes of 0.848·10^−3^, 1.74·10^−3^, 3.63·10^−3^ s^−1^ for stresses of 4.68, 9.16, and 18.7 Pa, respectively. These straight segments, indicating constant shear rates at constant shear stresses, correspond to liquids with viscosities of 5510, 5260, and 5150 Pas, respectively. When the stresses are dropped to zero, the material experiences a minor recoil over ~10 sec, indicating elastic properties, and the deformation remains steady afterwards. Therefore, the overall mechanical response of material corresponds to a viscoelastic fluid with some shear-thinning of viscosity that is typical for polymeric liquids. We note that the viscosity of the material resulting from B:C = 85:1 is ~1000 times greater than of the base component of Sylgard 184 (5.1 Pas according to the manufacturer), suggesting that the addition of the curing agent has caused substantial polymerization, which still has not lead to solidification. (D) Mechanical tests of the material obtained by mixing the components of Sylgard 184 at B:C = 100:1 and curing the mixture overnight in a forced convection oven at 80 °C. Using the shear rheometer, at time t = 0 a 191 μm thick layer of the material was subjected to a certain shear stress, at a time ~180 sec the shear stress was dropped to zero, and the shear strain of the layer was measured as a function of time. The values of the shear stress applied between 0 and 180 sec were 1.75 Pa (black dots), 4.85 Pa (orange dots), and 9.5 Pa (blue dots). The segments with the non-zero shear stress are well fitted by straight lines with slopes of 0.871·10^−3^, 2.43·10^−3^, 4.73·10^−3^ s^−1^ for stresses of 1.75, 4.85, and 9.5 Pa, respectively. These straight segments, indicating constant shear rates at constant shear stresses, correspond to liquids with viscosities of 2010, 2000, and 2000 Pas, respectively. After the stresses are dropped to zero, the shear strain remains constant (apart from a minor instrumental zero drift). Therefore, the overall mechanical response of the 100:1 mixture corresponds to a viscous Newtonian liquid (possible non-Newtonian properties are undetectable in our test) with viscosity ~400 times greater than the viscosity of the base component of Sylgard 184. We finally note that the plot in panels C and D, all representing liquids, are in a stark contrast with the plot in Fig. S1H, showing the results of a similar mechanical test performed on a soft solid gel with *E* = 0.44 kPa. In that last case, the deformation (shear strain) reaches a plateau within a short time after shear stress is increased and fully recoils after the shear stress is reduced to its previous value.

**Figure 8.**
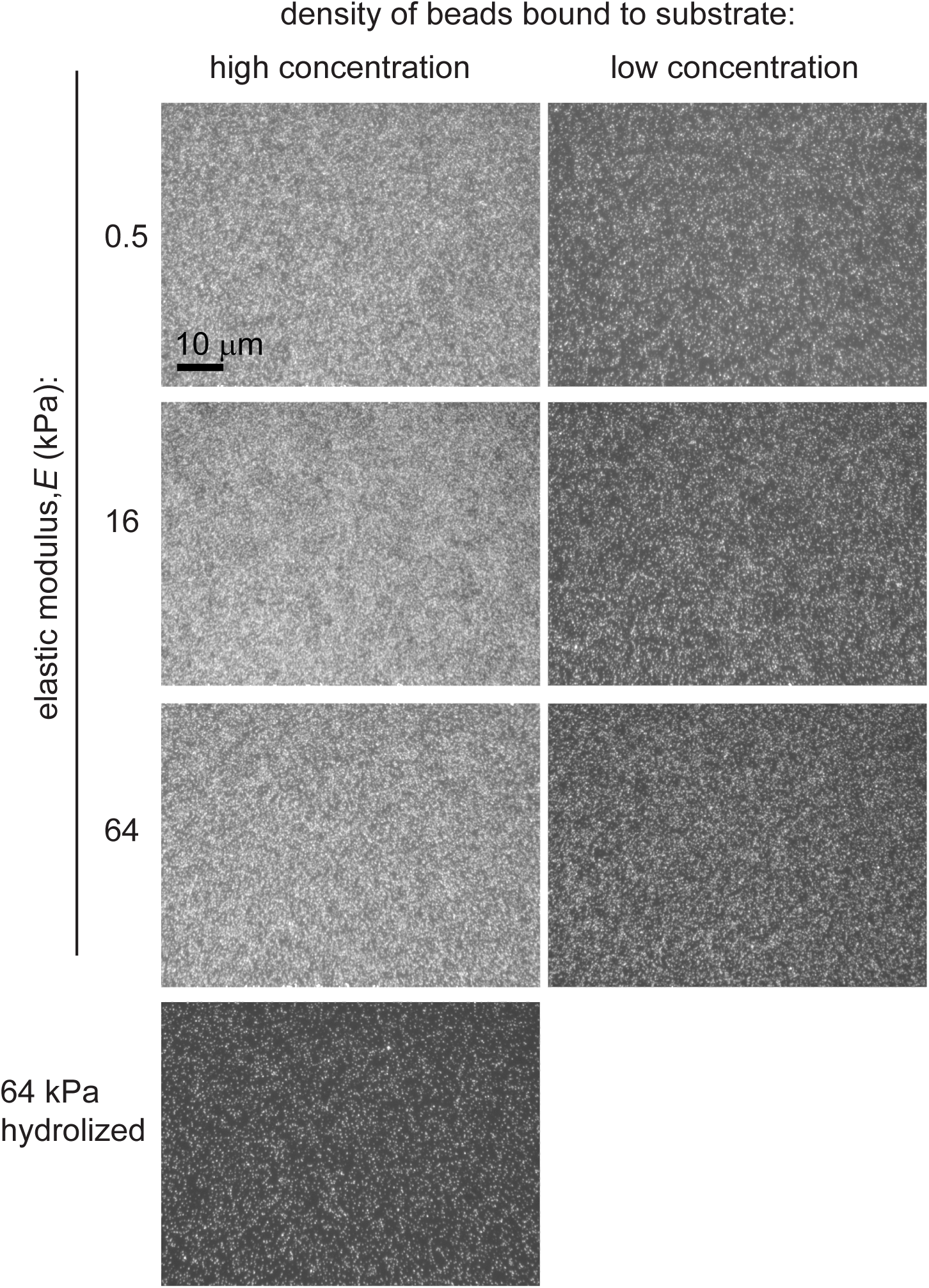
Binding of functionalized beads to silicone gels substrates. Fluorescence images of 40 nm beads (580nm excitation/605nm emission) on surfaces of silicone substrates with different elastic moduli after the substrates were incubated under bead suspensions for 30 min at room temperature (RT). The beads had amine groups on their surfaces. Two bead suspensions were used with the low-concentration suspension been 1/8 as concentrated as the high-concentration one. In a control experiment with a 64 kPa gel (*bottom left*), amine-reactive groups were hydrolyzed by incubation under 20 mM HEPES buffer (pH 8) for 30 min at RT before the high-concentration bead suspension was applied for 30 min at RT.

## Discussion

In their experiments, both Trappmann *et al.*^4^ and Wen *et al.*^3^ used silicone substrates prepared by mixing different proportions of the components B and C of Sylgard 184 elastomer (Dow Corning). The values of the elastic moduli, *E*, for Sylgard 184 mixtures with a range of B:C have been measured by several groups using different experimental techniques. These values are plotted as a function of B:C in Fig.7A. There is a good agreement between the values of *E* obtained from the extension tests on cylindrical samples by Cesa *et al.*^9^ (Fig. 7A, black triangles) and our measurements under shear using the specially built shear rheometer (Fig.7A, green circles). In addition, there is a good agreement between the results obtained with the shear rheometer and with our microfluidic tests for B:C from 20:1 to 30:1 and from 65:1 to 75:1. On the other hand, in the range of B:C from 40:1 to 55:1, the microfluidic tests measured *E* as much as 1.5-times lower (or corresponding to B:C ratios ~5 points greater) than the tests with the rheometer and the tests on cylindrical samples. The value of *E* for B:C = 50:1 from Trappmann *et al.*^4^ (Fig.7A, blue squares) is ~1.5 times greater than the values measured with the cylindrical samples and with the shear rheometer and the rest of the values of *E* in the interval of B:C from 20:1 to 50:1 from Trappmann *et al.*^4^ are 1.5-2 times greater than the values of *E* obtained from the measurements under extension and shear. These discrepancies can be due to variability of properties between different batches of Sylgard 184. Indeed, there is a considerable variability in the values E measured at different times (and hence, for different samples and batches) by the same groups (Fig.7A). It is worth noting, however, that the measurements using shearing of thin layers and extension of cylinders (or slabs) both rely on simple deformations and involve minimum assumptions about mechanical properties of the gels, and their results depend on the Poisson ratios of the gels (expected to be close to 0.5) only weakly. In contrast, the deformations produced by indenters (both flat and round) are of a relatively complex and sample thickness-dependent profiles, and the experimental readout may be more dependent on the Poisson ratio and on the details of interaction between the indenter and the surface of the material.

The data point for B:C = 100:1 from Trappmann *et al.*^4^ (Fig.7A, blue squares), with *E* = 0.1 kPa, does not appear to be inconsistent with the trends in the data obtained with the stretching of the cylindrical samples and with the shear rheometer and microfluidic technique (although an extrapolation of those two last sets of data to B:C = 100:1 would give a value of *E* an order of magnitude below 0.1 kPa). Nevertheless, we have never been able to make mixtures with B:C ≥ 80:1 to form solids. They always remained liquid even after multiple days of curing at an elevated temperature of 100°C (Fig.7B). Rheological tests on a B:C = 100:1 mixture cured overnight at 80°C (Fig.7D) indicated that it was a Newtonian liquid with a viscosity of 2000 Pas, which was ~400 greater than the viscosity of component B of Sylgard 184 (5.1 Pas according to the manufacturer). Thus, the addition of 1% component C lead to substantial polymerization (hence the increase in viscosity), but not curing. Rheological tests on a B:C = 85:1 mixture cured overnight at 80°C indicated that it was a viscoelastic liquid with a viscosity of 5500 Pas, some shear-thinning, and detectable elastic properties (detectable recoil after the stress is dropped; Fig.7C). Thus, as expected, the addition of a greater amount of component C (~1.17% vs. 1%) lead to increased polymerization (hence the increase in viscosity and detectable elasticity), but still did not produce a solid material (Fig.7C). It is instructive to compare the plots in Fig.7C and D, showing mechanical responses of liquids, with the plot in Fig.1H, showing the mechanical response of the 0.5 kPa SoftSubstrates gel, which is a solid. Whereas in the two first cases constant shear stresses lead to constant shear rates, and the samples remain deformed after the stresses are relieved, in the last case, after a stress is applied, the deformation rapidly reaches a plateau and, after the stress is relieved, the sample regains the shape it had before the application of the stress.

It is our understanding that the elastic modulus for the material with B:C = 100:1 was measured in the test presented in Fig.S1d in Ref.^4^, in which a layer of the material was indented with a 20 mm flat punch. The elastic modulus of 0.1 kPa was then calculated as a number proportional to the value of load force at time going to infinity. As far as we can judge from the plot in Fig.S1d in Ref.^4^, the value of the load force is <0.1 g (~5% of the peak value of load in the test) at the largest time plotted (~270 sec) and continues decaying with time. Therefore, the fact that the measured load force is different from zero (which would be expected from a liquid in a steady state) may be explained by a combination of an insufficient time of observation and limited accuracy of the experimental system (zero drift *etc*.). Thus, in our view, the results of the test in Fig.S1d in Ref. ^4^ do not clearly demonstrate that the sample with B:C = 100:1 is a solid.

It also did not escape our attention, that when keratinocytes were plated on substrate with B:C = 100:1, the substrate had a visible pattern of wrinkles on a scale of several μm (top left panel in Fig.1C in Ref.^4^). This pattern is reminiscent of the pattern of wrinkles that was produced by cells plated on the surface of liquid silicone with a thin hard crust, which was formed by flaming the silicone surface^10^ (and apparently resulted from the buckling instability of the crust under the action of the traction forces of the cells). Therefore, we suspect that a similar thin hard crust might have been formed on the surface of the B:C = 100:1 material in Ref.^4^, when it was treated with UV-light and reactive chemicals for the purpose of functionalization of the surface.

The above considerations suggest that the phenotypes of cells plated on the substrates with B:C = 100:1 in Ref.^4^ may reflect the responses of the cells to the unknown rigidity of the hard crust rather than to the stated elastic modulus of the material (0.1 kPa). Therefore, the phenotypes of cells plated on the substrate with B:C = 100:1 cannot be reliably qualified as phenotypes of cells plated on a soft gel. We note that in Ref. ^4^, the data on cells plated on the substrate with B:C = 100:1 are the only data on cells plated on either soft (*E* < 4 kPa) or medium rigidity (*E* = 8 −17 kPa) substrates. If the data on cells on the B:C = 100:1 substrate are disregarded, the only data remaining are on cells on hard substrates, with *E* ≥ 40 kPa (which may also have hard crusts on their surfaces). For PAA gels in this range of *E*, the dependence of cell phenotypes on the substrate rigidity is usually found to be weak or insignificant (cf. Fig.1 and Fig.3 in Ref.^4^). Hence, the similarity of phenotypes of cells plated on the silicone substrates used by Trappmann *et al.*^4^ does not imply that cells do not respond to rigidity of silicone gels or that cellular responses to rigidity of silicone gels are different from cellular responses to rigidity of PAA gels.

In their paper, Wen *et al.*^3^ state *“These results, in conjunction with cure ratio-independent stem cell spreading (Supplementary Fig.11B) and differentiation (Fig.4E), emphasize the shortcomings of PDMS as a model system to investigate stiffness-dependent behaviour over a relevant cell-sensing range.”* The phrase “these results” in the beginning of this sentence refers to the difficulty of functionalization of surfaces of silicone gels with reactive groups; we will address this subject later. As for the results shown in Fig.4E and Supplementary Fig.11B in Ref.^3^, they indicate that both MSC differentiation and ASC spreading are practically identical for cells plated on substrates obtained from mixing the B and C components of Sylgard 184 at 50:1, 75:1, and 100:1. We immediately note, however, that both 50:1 and 100:1 substrates were found to be more rigid than the PAA gel with *E* = 30 kPa (Fig.4D in Ref.^3^; the mechanical characterization for the 75:1 mixture was not presented). As argued above, for hard substrates, with *E* > 40 kPa, the dependence of cell phenotypes on the substrate rigidity is expected to be weak or insignificant. Therefore, the similarity of phenotypes of cells plated on the 50:1, 75:1, and 100:1 substrates (Fig.4e and Supplementary Fig.11B in Ref.^3^) is consistent with the mechanical properties of these substrates, as they are presented in Ref.^3^.

The experimental data obtained by us^11,12^ (Fig.1 and Fig.7A) other authors^9,13–15^ (Fig.7A) indicate that silicone gels can be made of soft and of medium rigidity. Here we presented results indicating that the rigidity of silicone gels affects the phenotypes of cells plated on them in the same way as the rigidity of PAA gels^4,3,2,16^ and micropost arrays ^5^.

A salient feature of the plot in Fig.7A is that, when converted to elastic moduli, *E*, the values of rigidity of the Sylgard 184 silicone gels with B:C = 50 and 100 measured by Wen *et al.*^3^ (*red squares*; they are reported as substrate spring constants) are visibly incompatible with the values and trends of elastic moduli measured by the other groups. Specifically, for B:C = 50, *E* is 20-50 times greater than the values obtained in the other studies and, for B:C = 100, which did not form a solid gel in our tests, *E* is on the same order as measured in the other studies for B:C = 36 – 45. We cannot know the exact reasons for this discrepancy. However, an obvious consideration is that the measurement technique used by Wen *et al.*^3^ relies upon the resistance of the substrate to the retraction of an AFM tip brought into contact with the surface of the substrate. Hence, results of measurements with this technique may specifically depend on the properties of the surface of the substrate (or a thin layer at its very top) and thus may not be universally related to the elastic properties of the bulk of the substrate. The relation between the resistance force and the elastic modulus of the substrate may not be a simple proportionality and may be different for silicone and PAA gels. If this is the case, our conversion of the spring constants measured by Wen *et al.*^3^ to elastic moduli is not legitimate and there is also no obvious way to perform such a conversion or unambiguously relate the results obtained by Wen *et al.*^3^ for cells on silicone substrates to the existing literature, in which the substrates were characterized by elastic moduli of their bulk^11,9,12–15^.

Fig.4E and Supplementary Fig.11B in Ref.^3^ show similar MSC differentiation and ASC spreading on the B:C = 50:1 and 75:1 substrates. The measurements by different groups (Fig.7A, all data except for red squares) suggest that these two substrates should be, respectively, of medium-to-high rigidity (13-40 kPa) and soft (<1 kPa) that would imply significant differences between spreading and differentiation for cells plated on these substrates. Notably, however, Wen *et al.*^3^ also found similar spreading of ASC and differentiation of MSC on the substrates with B:C = 100:1, whereas we have not been able to make mixtures with B:C ≥ 80:1 to solidify. Therefore, an explanation of the results in Fig.4e and Supplementary Fig.11B in Ref.^3^ could be that the 100:1 and 75:1 substrates (and possibly the 50:1 substrate) had hard crusts formed on their surfaces. The formation of the hard crusts would then also explain the high values of the substrate spring constants measured for these silicone substrates (Fig.4D in Ref.^3^).

Finally, we note that, when untreated, the surfaces of both PDMS (and other silicones) and PAA are chemically inert and do not spontaneously form covalent bonds with ECM proteins. Hence, whereas some ECM may bind to surfaces of both PDMS and PAA by passive absorption, for strong and reliable binding of ECM, the gel surfaces need to be functionalized with some reactive groups. The chemical structures of PDMS and PAA are essentially different, however, such that the functionalization of surfaces or PDMS and PAA gels requires different reagents. So, as noted by Wen *et al.*^3^, sulpho-SANPAH, a standard reagent for functionalization of PAA, cannot be expected to be efficient for PDMS. Strangely, Wen *et al.*^3^ stated that “PDMS substrates do not support protein tethering” because there is an insufficient linking of the ECM to silicone gel surface. We do not see sufficient evidences supporting this statement. The surface chemistry of PDMS gels can be readily modified with a variety of silanes with reactive groups^12,17^, and in this study we used silicone gels with surfaces functionalized with amino-reactive groups (Fig.8). Therefore, in our view, ability to functionalize their chemically inert surfaces for ECM binding makes silicone gel substrates a versatile tool to study cellular rigidity sensing.

## Materials and Methods

### Silicone gel substrates for cell culture

6-well plates with silicone gels on the well bottoms (SoftSubstrates) were obtained from MuWells (softsubstrates.com, San Diego, CA).

### Measurements of elastic moduli of silicone gels substrates

Elastic moduli of the silicone gels were measured by assessing the deformation of thin layers of gels under known shear stresses using a previously reported microfluidic technique^11^ (Fig. 1A-C), a newly developed gel rheometer^18^ (Fig.1D-F), and a modified version of the microfluidic technique (Fig.1G-I). In the first microfluidic technique, thin uniform layers of the silicone gels (30 - 65 μm thickness) used in the 0.5, 16, and 64 kPa 6-well plates were prepared on 50x35 mm #1.5 microscope cover glasses with 100 nm fluorescent beads deposited on the cover glass surfaces (special order from MuWells). The gel surfaces were coated with 40 nm red fluorescent beads (see below) and the thicknesses of the gel layers, *ξ*, were measured under a fluorescence microscope by focusing first on the beads on the surface of the coverglass and next on the beads on the surface of the gel. A specially designed microfluidic device was bonded to the surface of each gel substrate. The device was perfused with a 80:20 mixture of glycerol and water (by weight) by applying controlled differential pressures, Δ*P*, between the inlet and outlet of the device. The displacements of the beads on the gel surface, Δ*x*, near the middle of the bottom of a 2 mm long, 2 mm wide and *h* = 165 μm deep channel of the device were measured under a fluorescence microscope at different hydrodynamic shear stresses, *τ*. The values of *τ* were calculated from Δ*P* using a prior calibration of the device. (The calibration was performed by perfusing the device with a 80:20 glycerol:water mixture with viscosity *η* = 0.060 Pas, which was seeded with 4.6 μm green fluorescent beads. The flow velocity at the mid-plane of the channel, *v*, was assessed at different Δ*P* by visualizing the bead streak lines, and the shear stress was calculated as τ = 6*ηv* / *h*.) The displacement of beads on the gels surface, Δ*x*, was used to calculate the substrate shear strain, *γ* = Δ*x* / *ξ*. The dependences of *γ* on *τ* were measured for all three gel types (Fig.1A-C). There was no detectable hysteresis or creep (slow growth of strain under constant stress) in any of the tests and all three dependencies of *γ* on *τ* were well fitted by straight lines passing through the origin, such that all three types of gels behaved as proper solids. The slopes of the straight line fits were used to calculate the shear moduli of the gels, *G* = *τ* / *γ*. The elastic moduli (Young’s moduli) of the gels were calculated as *E* = *G* / 3 = *τ* / 3*γ* (assuming a Poisson ratio of 0.5)^11^. The values of *E* were found at 0.61, 20, and 62 kPa for the 0.5, 16, and 64 kPa gels respectively.

With the newly developed gel rheometer, the deformations of 20x30 mm rectangular gel layers with thickness from 1.35 to 1.54 mm were tested under centrifugal force ^18^, and the dependences of the average shear strain, *γ*, on the average shear stress, *τ*, were measured (Fig.1D-F). The dependencies were fitted again by straight line passing through the origin, and their slopes were used to calculate the shear moduli, *G* = *τ* / *γ*, and the elastic moduli *E* = *G* / 3 = *τ* / 3*γ* of the gels (assuming again a Poisson ratio of 0.5)^11^. The values of *E* were found at 0.40, 17, and 65 kPa for the 0.5, 16, and 64 kPa gels respectively (Fig.1D-F). The values of the elastic moduli measured by the two techniques for the gels with the nominal elastic moduli of 16 and 64 kPa are in good agreement with each other and with the nominal values (20 and 17 kPa vs. 16 kPa; 62 and 65 kPa vs. 64 kPa). The discrepancy between the values of *E* measured with the two techniques (0.61 and 0.40 kPa) and the stated value of *E* for the 0.5 kPa gel is greater and is likely due to batch-to-batch variability (all gels samples were from different batches). Nevertheless, the uncertainties in the elastic moduli of the 0.5, 16, and 64 kPa gels, as estimated from our mechanical measurements, do not affect the major conclusions from any of our experiments on cells plated on these gels (cf. Fig.2, Fig.3-6).

To check for possible variation of the effective elastic moduli of the silicone gels on scales <30 μm, we used a modified version of the microfluidic technique to test 18, 6.3, and 2.4 μm thick layers of the SoftSubstrates gel with nominal *E* = 2.0 kPa (special order from MuWells). As before, there was a low concentration of fluorescent beads (40nm, red fluorescent) on the coverglass-gel interface and a higher concentration of the same beads covalently bonded to the gel surface. The thicknesses of the gels were assessed by taking stacks of fluorescence images on a confocal microscope with a precise z-axis encoder and finding the z-axis positions at which the beads on the interface and the gel surface were maximally in-focus. The modifications of the microfluidic technique were two-fold. First, the dimensions of the test chamber were reduced ~3-fold (as appropriate for the relatively thing gel layers tested), to ~600x600x60 μm. Second, instead of the 80:20 glycerol:water mixture, the working fluid was Helium, thus enabling faster measurements without the risk of spill. (The correspondence between *τ* at the gel surface at the bottom of the test chamber and Δ*P* between the inlet and outlet of the device was calibrated by perfusing the device with a suspension of fluorescent beads in water and measuring the streak lines of the beads at various Δ*P*; the correspondence was theoretically predicted to be the same for Helium as for water.) For all three gel thicknesses, the dependences of the shear strain, *γ* = Δ*x* / *ξ*, on *τ* were well-fitted by zero-crossing straight lines (Fig. 1G). The values of *E* calculated from the dependencies, 1.70, 1.78, and 1.65 kPa, respectively, for the 18, 6.3, and 2.4 μm thick gels, were consistent with each other, with the value of 1.9 kPa from our measurement on a 1.05 mm thick sample using the shear rheometer, and with the nominal value of 2 kPa. The results of this test indicated that the effective elastic modulus of the gels used in our study is uniform down to a subcellular scale of 2.4 μm.

Finally, we used the rheometer to measure changes in shear strains of gel layers, *γ* = Δ*x* / *ξ*, upon step-wise changes in the shear stress in order to estimate relaxation times of the gels (Fig. 1H-J). The relaxation time of a gel corresponds to a temporal scale at which the effective viscosity of the gel is as important as its elasticity. Processes with time scales shorter than the relaxation time are expected to be viscosity-dominated, whereas processes with time scales longer than the relaxation time are elasticity-dominated. Based on results of these tests, the relaxation times of the 0.5, 16, and 64 kPa gels (actual values of *E* measured at 0.44, 18, and 64.5 kPa, respectively) were estimated at <4, <1, and <1 sec, respectively. A relaxation time of 4 sec would certainly be a major factor for some rapid cellular processes sensitive to the mechanics of the substrate, such as changes in contractility of cardiomyocytes. On the other hand, changes in cellular adhesions and contractility during the spreading of cells and differentiation of stem cells occur on substantially longer time scales (tens of second)^19^, rendering the relaxation time of 4 sec too short to make the viscous properties of the 0.5 kPa gel relevant for these cellular processes. (The relaxation times of the 16 and 64 kPa gels are even shorter, making their viscosities even less relevant.)

### Assaying density of amine-binding sites on silicone gel substrates

To assess the density of amine-binding sites on the silicone gel substrates, we used 40 nm red fluorescent beads functionalized with amine groups. The beads were prepared by incubating 40 nm carboxylated fluorescent beads (580nm excitation/605nm emission; F-8793 by Life Technologies) with silane-PEG-amine (from Laysan-Bio, Arab AL). The bead suspensions were used at two dilutions, 1:20,000 (high concentration) and 1:160,000 (low concentration), in 20 mM HEPES buffer (pH 8). To assay the binding of beads to the silicone gel substrates, bead suspensions were incubated in wells of the silicone gel-bottom 6-well plates for 30 min at room temperature (RT). After the incubation, the wells were thoroughly rinsed with water and the fluorescent beads attached to silicone gel surfaces at the well bottoms were photographed under a fluorescence microscope. The assay was performed in 6-well plates with gels of all three elastic moduli and with both low-concentration and high-concentration bead suspensions (Fig.5). The bead density on the gel substrates was found to be consistent between substrates of all three elastic moduli for both high-concentration and low-concentration bead suspensions.

To test for the specificity of the binding of beads to the gel substrates, a 64 kPa gel on a well bottom was incubated for 30 min under a pH 8.5 HEPES buffer and then for 30 min under the high-concentration bead suspension. The pre-incubation under pH 8.5 HEPES was expected to result in hydrolysis of the amino-reactive groups on the gel surface. The density of fluorescent beads on the gel surface was found to be greatly reduced as compared with the bead density obtained with the same high-concentration suspension without the pH 8.5 HEPES pre-incubation and was also markedly lower than the density obtained with the low-concentration bead suspension without the pH 8.5 HEPES pre-incubation.

Taken together, these results indicate that the surfaces of the silicone gels of all three elastic moduli used in our study are indeed functionalized with amino-reactive groups and the effective site density of these groups is similar for gels of all three rigidities. Therefore, a 6-well plate used in the study is filled with a solution of ECM proteins, the protein molecules are expected to covalently bind to the gels surfaces through their amino groups, with the covalent binding being equally efficient for gels of all three elastic moduli.

### Stem cell differentiation

Human mesenchymal stem cells (hMSCs) of early passages (P0) were obtained from Stemedica (San Diego, CA). Silicone gels substrates were coated with 1.6 μg/ml solution of collagen I (Advanced Biomatrix, San Diego, CA) in pH 7.4 PBS for 30 min at 37 °C. hMSCs were seeded into the 6-well plates at 600 cells/ cm^2^ in 2 mL of 7.5% BGS (EquaFETAL^®^, AtlasBIOLOGICALS) hMSC growth media (Stemedica) and grown in humidified oxygen-controlled 37 °C incubator with 5% O_2_ and 5% CO_2_. Cells were allowed to reach ~75% confluence before a differentiation medium (ThermoFisher) was applied to induce either adipogenesis or osteogenesis. After 7 days, the differentiation medium was refreshed, and after 14 days cell were examined to assess their differentiation. Adiposeness was assessed using AdipoRed (ThermoFisher), according to a protocol recommended by the manufacturer with the following modifications: prior to the addition of AdipoRed, all cells from the wells of a 6-well plate were harvested by trypsinization, washed once in pH7.4 PBS, resuspended in 1.2 ml of PBS and transferred into a 96-well plate (200 uL of cell suspension per well); AdipoRed was added to each well of the 96-well plate, incubated for 20 min at RT, and the intensity of staining was measured using a fluorimeter (FLX800, Biotech Instruments Inc). Osteoblasts were stained with Alizarin Red and imaged using Evos FL cell imaging system (Advanced Microscopy Group, Mill Creek, WA), with the level of osteogenesis assessed as previously described^20^. Alternately, osteogenesis was assayed as described in Ref. ^4^. Briefly, hMSCs were seeded at a density 2,000 cells/cm^2^, cultured for 1 hour, and the differentiation medium was applied. Cells were assayed for ALP activity after 7 days.

### Cell spreading assay

To measure hMSC spreading areas, hMSCs were plated onto silicone gel substrates in 6-well plates and cultured for 24 hours as described above in “stem cell differentiation”. Cells were then fixed with 3.7% formaldehyde in PBS, permeabilized with 0.5% Triton X-100 in PBS at RT for 10 min, and washed three times with PBS. The fixed cells were incubated with phalloidin-conjugated rhodamine (Molecular Probes) for 45 min at RT and washed three times with PBS. Next, cells were photographed under a fluorescence microscope. Mouse primary keratinocytes and mouse embryonic fibroblasts (MEFs) were plated on ~30 μm layers of the 0.5, 16, and 64 kPa silicone gels on #1.5 microscope cover glasses (special order from MuWells, making it possible to measure their spreading areas under fluorescence microscope with improved resolution. Silicone gel surfaces were coated with fibronectin (ThermoFisher) by incubation under a 20 μg/ml solution of fibronectin in pH 7.4 PBS for 30 min at RT. Keratinocytes and MEFs were plated on fibronectin-coated silicone gels and incubated in DMEM supplemented with 1% (v/v) BSA for 45 min at 37 °C,5% CO_2_. Cells were fixed and stained with phalloidin as described above. Next, the cover glasses were mounted on microscope slides with a mounting solution (ProLong^®^ Gold antifade reagent; Invitrogen) and cells were photographed under a fluorescence microscope. The micrographs were digitally processed and cell spreading areas were quantified using a code in MATLAB (MathWorks, Natick, MA). For each substrate elastic modulus and each cell type, the spreading areas were measured for 75 cells in randomly selected areas of the substrate (Fig.4,5).

### Preparation of cell lysates and Western blotting

Keratinocytes and MEFs were plated on fibronectin-coated silicone gel substrates in the 6-well plates and incubated for 45 min at 37°C, 5% CO2. Whole cell lysates were prepared using modified radioimmune precipitation assay buffer (50 mM Tris, pH 7.5, 150 mM NaCl, 50 mM NaF, 1 mM sodium pyrophosphate, 0.1% sodium deoxycholate, 1% Nonidet P-40, protease inhibitors cocktail and 1% CHAPS). Lysate protein was quantified using the bicinchoninic acid (BCA) method (ThermoFisher Scientific), normalized, and used in Western blots analysis. The primary antibodies for Western blots were against Y^576^FAK (ThermoFisher), FAK (Cell Signaling), and α-tubulin (Sigma) (Fig.5).

